# Brain dynamics of memory encoding for simple versus complex musical sequences

**DOI:** 10.64898/2026.08.28.747779

**Authors:** Gemma Fernández-Rubio, Leonardo Bonetti, Morten L. Kringelbach, Peter Vuust, Elvira Brattico

## Abstract

Memory encoding is the foundational process by which the brain transforms sensory input into lasting representations. While the neural mechanisms of auditory memory have been extensively studied, how musical complexity modulates the neural activity during memory encoding remains poorly understood. Here, we used magnetoencephalography (MEG) to investigate the encoding of simple (tonal) versus complex (atonal) musical melodies in 67 participants. Behaviorally, the latter melodies were consistently rated as more complex and associated with lower recognition accuracy across three testing sessions (same day, one day later, and ten days after the encoding task). At the neural level, source-localized analyses revealed distinct spatiotemporal dynamics: simple melodies elicited stronger activity in auditory cortices (left and right Heschl’s gyrus) and cingulate regions (medial and anterior cingulate gyrus), while complex melodies recruited the left hippocampus more extensively across multiple tones. These findings demonstrate that musical complexity shapes neural encoding processes from the outset, with tonal sequences benefiting from efficient sensory processing and atonal sequences requiring greater memory-related recruitment. Our study provides novel insights into how the human brain encodes complex auditory information, providing a framework for understanding the neural basis of memory formation for temporally structured stimuli.

## INTRODUCTION

Memory and cognition rely on the brain’s capacity to extract, predict, and integrate temporal patterns from the environment. This ability is essential for survival, enabling us to anticipate events, recognize familiar stimuli, and form lasting memories. Auditory sequences, such as music, provide a simple, unique, and ecologically valid model for studying these processes, as their meaning emerges from the precise temporal organization of individual sound elements over time (Brattico & Delussi, 2024; Peretz & Zatorre, 2005). Unlike static visual stimuli, auditory information unfolds dynamically, requiring the brain to continuously integrate incoming sensory input with existing memory representations to support perception, prediction, and recognition (King et al., 2016; King & Wyart, 2021; Koelsch et al., 2019; Marti & Dehaene, 2017; Zatorre, 2003).

Our previous investigations have begun to unravel the neural mechanisms underlying auditory memory recognition. In a series of magnetoencephalography (MEG) studies, we demonstrated that recognition of previously learned auditory sequences relies on a widespread cortico-subcortical network comprising both sensory and memory-related regions (Bonetti et al., 2023; Bonetti et al., 2024a; Bonetti et al., 2024b; Bonetti et al., 2024c; Fernández-Rubio et al., 2022a; Fernández-Rubio et al., 2022b; Serra et al., 2023). Specifically, we showed that recognition of tonal (i.e., simple) versus atonal (i.e., complex) musical sequences recruits distinct neural pathways (Fernández-Rubio et al., 2022b). Tonal sequences, which are prototypical of Western popular music, engaged hippocampal and cingulate areas, while atonal sequences, which is the direct opposite of tonal music, primarily activated the auditory network. Previous studies collectively demonstrate that tonal and atonal music differ primarily in the encoding schemas they employ. Tonal music leverages well-learned pitch hierarchies to enable categorical coding, structural reduction, efficient chunking, and precise expectations. Atonal music, by contrast, offers fewer universally shared pitch-stability cues, placing greater demands on auditory rehearsal, attention, segmentation, and associative memory. These differences initially make atonal music more difficult to remember, predict, and emotionally interpret, particularly for listeners without relevant expertise (Daynes, 2011; Mencke et al., 2018; Mencke et al., 2021; Mencke et al., 2022; Ockelford & Sergeant, 2013; Proverbio et al., 2015; Serra et al., 2012; Vuvan et al., 2014; Wald-Fuhrmann et al., 2026). Our findings support the literature by showing that musical complexity qualitatively alters the neural pathways of recognition memory, with simpler, more predictable auditory information relying more heavily on memory-related regions, whereas more complex, less predictable sequences depend on sensory processing areas. These neural differences align with behavioral evidence that tonal and atonal sequences are encoded differently, with tonal music benefiting from categorical coding strategies that atonal music cannot support (Mikumo, 1992). Furthermore, in a subsequent study, we uncovered the hierarchical brain organization during recognition of tonal musical sequences, characterized by feedforward connections from auditory cortices to hippocampus and cingulate gyrus, simultaneous with feedback connections operating in the opposite direction (Bonetti et al., 2024b). These findings provided evidence of hierarchical brain mechanisms during memory and predictive processing of auditory sequences.

While these studies have significantly advanced our understanding of auditory memory recognition, the neural dynamics underlying memory encoding, the initial stage of memory formation, remain poorly understood, particularly with respect to how musical complexity modulates this process. Memory encoding represents the critical first step in transforming sensory input into durable memory representations, yet the spatiotemporal brain mechanisms supporting the encoding of complex versus simple auditory sequences have not been systematically investigated.

Here, we address this gap by examining the neural activity during the encoding of tonal and atonal musical melodies. We employed a musical memory paradigm in which participants actively memorized simple tonal and complex atonal melodies while their brain activity was recorded with MEG. Based on our previous findings (Fernández-Rubio et al., 2022b), we hypothesized that the encoding of complex sequences would elicit distinct neural patterns compared to simple sequences, reflecting the increased cognitive demands and reduced predictability of processing atonal music. Our results reveal significant differences in brain activity between the encoding of simple and complex melodies, particularly in auditory and memory-related regions, and provide novel insights into how musical complexity shapes the neural mechanisms of memory formation.

## RESULTS

### Overview of experimental design

We recorded magnetoencephalography (MEG) data from 67 healthy participants while they performed a memory encoding task. Participants were presented with six musical melodies of varying complexity matched for rhythm, timbre, tempo, and meter (**Figure 1a**). During the MEG session, participants either passively listened to or actively memorized each melody in a counterbalanced design. For all analyses reported here, we focused exclusively on the “memorize” trials, excluding the passive listening data to isolate memory encoding processes. Following MEG acquisition, participants completed an old/new recognition task at three time points (same day, one day later, and ten days later) to assess memory retention (**Figure 1b**) and rated the perceived complexity of each melody. Source reconstruction was performed using a single-shell forward model and beamforming algorithm, yielding time series for 3559 brain voxels (**Figure 1c**). Building on our previous studies of musical memory recognition (Bonetti et al., 2024b; Bonetti et al., 2024c; Fernández-Rubio et al., 2022b), we focused our analyses on six anatomically defined regions of interest (left and right Heschl’s gyrus, left and right hippocampus, medial cingulate gyrus, and anterior cingulate gyrus) and estimated differences in neural activity between simple and complex melodies independently for each region (**Figure 1d** and **1e**).

**Figure 1.**
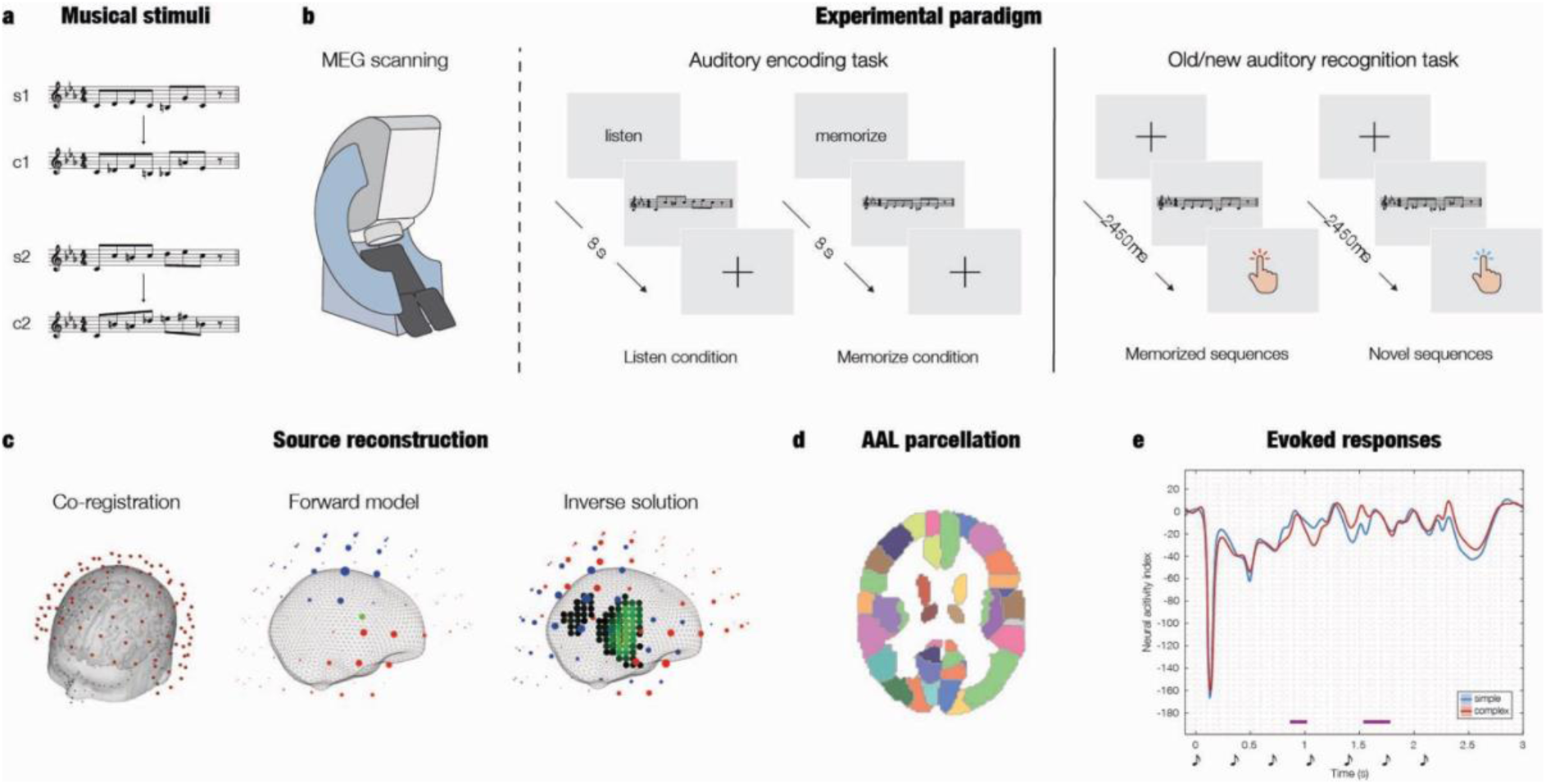
Experimental design and data analysis. **a.** The stimulus set comprised two tonal (s1, s2), two semi-tonal (m1, m2), and two atonal melodies (c1, c2), matched for rhythm, timbre, tempo, and meter. The tonal and atonal melodies used for further analyses are depicted. **b.** During magnetoencephalography (MEG) recordings, participants either passively listened to or actively memorized the six melodies. Half of the participants memorized melodies s1, m1, and c1 while passively listening to s2, m2, and c2; the other half did the opposite. After the MEG session, participants performed an old/new recognition task. They classified original (memorized) and modified (novel) versions of the melodies using one of two response keys. The task was administered on the same day as the MEG recordings, as well as one and ten days later. **c.** The MEG data from “memorize” trials was co-registered with the individual anatomical magnetic resonance imaging (MRI) data. Source reconstruction was computed using a single-shell forward model and beamforming algorithm as inverse solution, resulting in 3559 brain voxels. **d.** Brain voxels corresponding to six regions of interest (ROIs) using the automated anatomical labelling (AAL) parcellation method. **e.** The time series for each ROI and experimental condition were computed, and differences in neural activity between tonal (i.e., simple) and atonal (i.e., complex) melodies were estimated.

### Perceived complexity of melodies and response accuracy during recognition task

We conducted a Wilcoxon rank-sum test to compare the complexity ratings of the two melody types. The results revealed a statistically significant difference in ratings between the two conditions (W = 10,986, *p* < 0.001) (**Figure 2a**). This indicates that participants consistently rated atonal melodies (hereafter referred to as “complex”) as significantly more complex than tonal melodies (hereafter referred to as “simple”).

**Figure 2.**
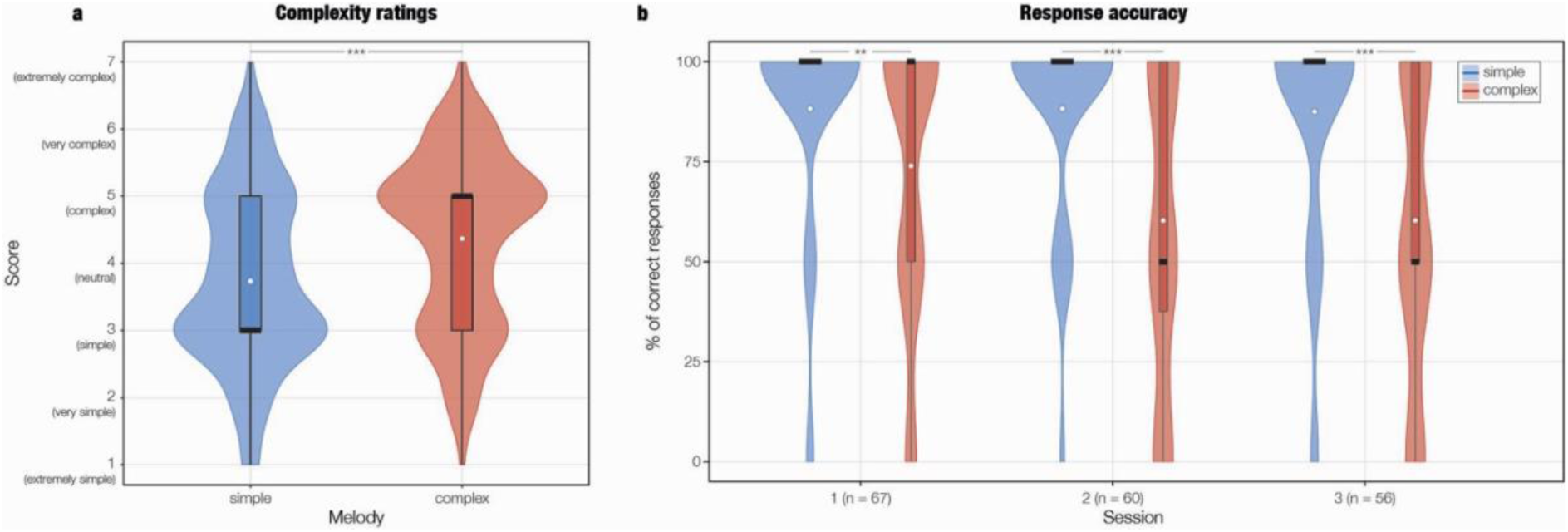
Differences in perceived complexity and response accuracy between simple and complex melodies. **a.** Violin and box plots show the distribution of perceived complexity ratings for tonal (“simple”) and atonal (“complex”) melodies on a 7-point Likert scale. White dots represent the mean response for each condition. A Wilcoxon rank-sum test confirmed a statistically significant difference in ratings (p < 0.001), with atonal melodies consistently rated as more complex than tonal melodies. **b.** Violin and box plots depict the effect of melody complexity on response accuracy in the old/new recognition task across three sessions (#1: same day as encoding; #2: one day later; #3: 10 days later). White dots represent the mean proportion of correct responses for each condition and timepoint. A generalized linear mixed-effects model revealed a significant main effect of melody complexity, with complex melodies associated with lower accuracy compared to simple melodies. No significant effects of time or complexity x time interaction were observed.

A generalized linear mixed-effects model (GLMM) with a binomial distribution and logit link function was fitted to assess the effect of melody complexity and time on response accuracy in the old/new recognition task. We found a significant main effect of melody complexity (β = –1.07, SE = .34, z = –3.14, *p* = .002), indicating that complex melodies were associated with lower response accuracy compared to simple melodies after controlling for time and participant variability. Post-hoc analyses further revealed that simple melodies were consistently associated with higher log-odds of correct responses than complex melodies across all sessions. This effect was most pronounced in session 2 (estimate = 1.81, p < .0001) (**Figure 2b**). There were no significant main effects of time (session 2: β = –0.06, SE = 0.40, *p* = .89; session 3: β = –0.34, SE = 0.39, *p* = .38), indicating that response accuracy did not change significantly across sessions. Additionally, the interaction between time and melody complexity was not significant (session 2 × complex: β = –0.74, SE = 0.49, *p* = .13; session 3 × complex: β = –0.44, SE = 0.48, *p* = .36), suggesting that the effect of melody complexity on response accuracy remained consistent across sessions.

### Brain activity during auditory encoding task

The timeseries of source-localized brain activity of “memorize” trials was contrasted between simple and complex melodies at six ROIs: the left and right Heschl’s gyrus (LHG, RHG), left and right hippocampus (LRP, RHP), medial cingulate gyrus (MCG), and anterior cingulate gyrus (ACG) (**Figure 3**).

**Figure 3.**
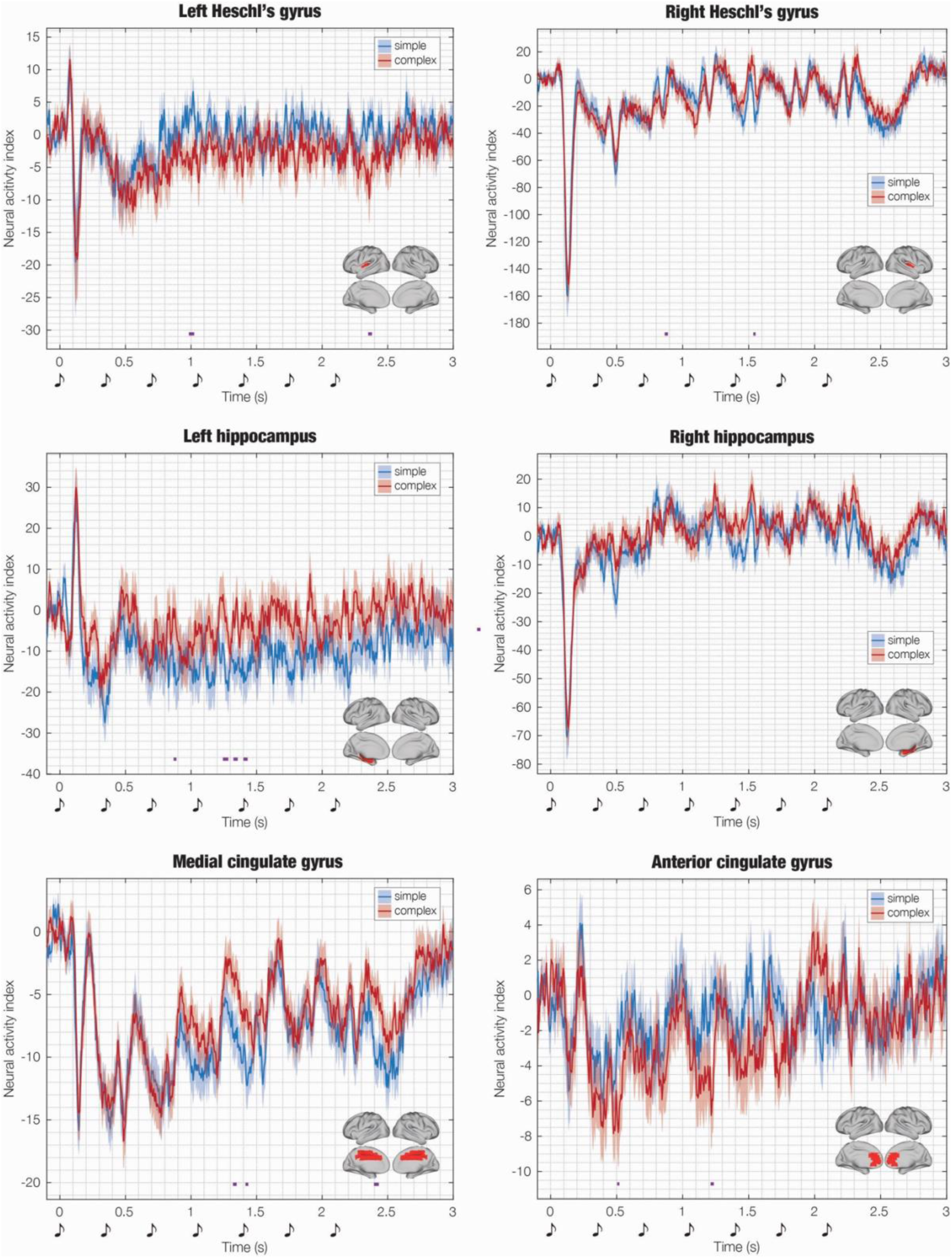
Contrast between encoding of simple versus complex melodies in source-localized brain activity. Time series data for six selected automated anatomical labelling (AAL) regions of interest (ROIs) are shown, comparing “memorize” trials of simple versus complex melodies. Purple lines indicate the temporal extent of significant differences between conditions, identified using cluster-based permutation testing (α = 0.05, 1000 permutations). Significant clusters were determined by comparing cluster sizes in the original data to a reference distribution derived from permuted data, with clusters deemed significant if their sizes exceeded those observed in 99.9% of permutations. Shaded areas represent standard errors. Musical sketches mark the onset of melody tones, and brain templates illustrate the spatial extent of the selected ROIs.

Overall, the contrasts revealed stronger activity in LHG, RHG, MCG, and ACG for simple melodies, particularly when the third and fourth tones of the melodies were presented (1–1.5 s), while the opposite effect was found in LHP. Four clusters of significant activity were found in LHP, denoting stronger activity for complex versus simple melodies between the third and sixth tones of the melodies. No significant differences between conditions were observed in RHP. Full results are reported in **Table S1**.

## DISCUSSION

This study provides novel insights into the neural mechanisms underlying the encoding of auditory sequences with varying degrees of complexity, extending our previous work on auditory memory recognition to the critical initial stage of memory formation. Using magnetoencephalography (MEG), we investigated how the brain encodes simple (tonal) versus complex (atonal) musical melodies, revealing distinct spatiotemporal dynamics that reflect the cognitive demands of processing predictable versus unpredictable auditory information. Our findings demonstrate that stimulus complexity not only modulates recognition memory, as we previously reported, but also shapes the neural mechanisms of memory encoding from the outset.

Our behavioral results confirm and extend previous observations regarding the processing of tonal and atonal music. Participants consistently rated atonal melodies as more complex than tonal ones, aligning with a substantial body of research demonstrating that atonal music is more challenging to process, less predictable, and less appreciated by non-expert listeners (Mencke et al., 2021; Mencke et al., 2022; Ockelford & Sergeant, 2013; Vuvan et al., 2014). Furthermore, our old/new recognition task revealed that complex melodies were associated with lower accuracy across all testing sessions, with the most pronounced differences observed one day after encoding. The absence of a significant time effect indicates that the difficulty in recognizing complex melodies persists across different retention intervals, highlighting the enduring impact of stimulus complexity on memory performance.

At the neural level, our source-localized MEG analyses reveal significant and region-specific differences in brain activity during the encoding of simple and complex melodies. We observed stronger activity in auditory cortices (left and right Heschl’s gyrus) for simple melodies, particularly during the processing of the third and fourth tones of the sequences (approximately 1–1.5 seconds after melody onset). This pattern suggests that predictable, tonal sequences benefit from more efficient sensory processing in auditory regions, potentially due to their adherence to familiar tonal hierarchies. Additionally, we observed stronger activity in the medial and anterior cingulate gyrus for simple melodies, regions associated with evaluative processes, suggesting that these areas may play a role in assessing the predictability of auditory input during encoding. Conversely, we found that the left hippocampus exhibited stronger activity for complex melodies across multiple tones, indicating that less predictable, atonal sequences recruit memory-related regions to a greater extent during encoding. Tonal music is organized around an internalized system of pitch relationships where tones vary in stability and structural importance. These tonal hierarchies support the perceptual organization of musical events, expectation formation, and the integration of individual notes into larger phrases and structures. Tonality can thus be understood as a learned cognitive schema that reduces uncertainty by enabling listeners to relate incoming events to stable reference pitches and anticipate likely continuations (Krumhansl & Cuddy, 2010). In contrast, atonal music is not perceived through the same strict event hierarchy. This does not imply atonal music is cognitively unstructured; rather, its representation may rely more on associative relationships, local similarities, recurring motives, and segmentation cues than on the pitch stability characteristic of tonality (Dibben, 1994; Imberty, 1993).

These results build upon and extend our previous investigations of auditory memory. In our earlier study, we demonstrated that the recognition of tonal versus atonal sequences engages distinct neural pathways, with tonal sequences recruiting hippocampal and cingulate areas and atonal sequences primarily activating auditory processing regions (Fernández-Rubio et al., 2022b). The current findings complement this work by revealing that similar distinctions emerge during the encoding phase, suggesting that the neural pathways supporting memory formation are modulated by stimulus complexity from the very beginning. The switch in hippocampal and auditory engagement between encoding and recognition reflects stage-specific cognitive demands. Tonal music’s predictability enables efficient sensory encoding in auditory and cingulate regions, while recognition requires hippocampal and cingulate retrieval to match stored representations. The involvement of the cingulate gyrus in both encoding and recognition further underscores its central role in auditory memory processes, potentially serving as a hub for integrating sensory input with memory representations and evaluative processes (Oane et al., 2023; Rolls, 2019). Conversely, atonal music’s reduced predictability necessitates hippocampal involvement for encoding but allows direct sensory comparison in auditory regions during recognition, as pattern matching is less effective for less structured material. Importantly, the experimental design of our previous study employed musical tones with a duration of 250 ms, which did not permit disentangling the overlapping activity of neighboring tones. As a result, the neural data had to be filtered in narrow frequency bands and averaged over each tone to make observations on the developing neural activity. The current study addressed this limitation by increasing the duration of the musical tones to 350 ms, a modification that revealed the distinct contributions of each tone forming the sequences to the broadband neural activity.

The observed differences in neural activity between simple and complex melodies can be interpreted within the predictive coding framework. According to predictive coding theory, the brain constantly generates predictions about incoming sensory input and updates its internal models to minimize prediction errors (Friston, 2010, 2012; Vuust et al., 2022). Tonal music, with its predictable structure and adherence to Western tonal hierarchies, may generate fewer prediction errors during encoding, allowing for more efficient processing in auditory regions (Krumhansl & Cuddy, 2010). In contrast, atonal music, characterized by the absence of a tonal center and increased unpredictability, recruits memory-related regions such as the hippocampus to encode the less predictable auditory information (Mencke et al., 2018; Mencke et al., 2021; Ockelford & Sergeant, 2013; Vuvan et al., 2014). This interpretation is consistent with our behavioral findings, as the increased cognitive demands of processing complex melodies are reflected in lower recognition accuracy. Moreover, it aligns with previous research demonstrating that prediction errors during encoding processes are associated with increased activity in the hippocampus (Aitken & Kok, 2022; Axmacher et al., 2010; Kumaran & Maguire, 2006).

While this study provides valuable insights into the neural mechanisms of auditory memory encoding, several limitations should be acknowledged. First, our stimulus set, while carefully controlled for rhythm, timbre, tempo, and meter, consisted of relatively short melodies, which may not fully capture the complexity of naturalistic musical listening. Future studies could employ longer, more ecologically valid musical stimuli to better approximate real-world experiences. Second, our focus on encoding precluded direct comparison with the recognition phase, as participants performed the recognition task after the MEG session. Future studies could address this by recording neural activity during both encoding and recognition to examine the continuity of memory processes and the transformation of neural representations from encoding to retrieval.

In conclusion, this study reveals that the encoding of simple versus complex auditory sequences relies on distinct neural pathways, with predictable, tonal melodies engaging auditory and evaluative regions and less predictable, atonal melodies recruiting memory-related areas. These findings extend our understanding of the spatiotemporal dynamics of auditory memory and highlight the importance of stimulus complexity in shaping neural mechanisms from the initial stages of memory formation. Our results demonstrate that the qualitative differences in neural processing observed during recognition are already present during encoding, suggesting that stimulus complexity modulates memory processes from the very beginning. Future work should continue to explore the hierarchical brain organization during both encoding and recognition and the generalizability of these findings to other types of stimuli. By doing so, we can gain a more comprehensive understanding of how the brain transforms sensory input into lasting memories and how this process is shaped by the complexity of the information we encounter in our daily lives.

## METHODS

### Sample

We collected magnetoencephalography (MEG) data from 71 healthy volunteers. Four individuals were excluded due to excessive noise or corrupted neural data, yielding a final sample of 67 participants (47 female; mean age = 25.88 years, SD = 6.91). Participants had heterogeneous musical backgrounds, with an average score of 23.28 ± 10.77 (maximum = 49) on the Musical Training subscale of the Goldsmiths Musical Sophistication Index (Müllensiefen et al., 2014). Musical expertise was not a recruitment criterion for this study.

All participants provided written informed consent and received modest financial compensation. The study was approved by the Institutional Review Board of Aarhus University (DNC-IRB-2022-011) and conducted in accordance with the Declaration of Helsinki – Ethical Principles for Medical Research.

### Experimental stimuli and design

The stimulus set comprised two tonal melodies (s1, s2), two semi-tonal melodies (m1, m2), and two atonal melodies (c1, c2). These were composed by GFR and matched for rhythm, timbre, tempo, and meter (**Figure 1a**). Each melody consisted of seven tones played on a piano, always beginning with middle C (261 Hz), arranged in an isochronous rhythmic pattern with 350-ms tone durations (**Figure S1**). MIDI versions of the melodies were created in MuseScore (v3.6.2) and presented using PsychoPy (v3.0).

A memory-encoding task was administered during MEG acquisition (**Figure 1b**). On each trial, participants were first shown the cue word “listen” or “memorize” for 1.5 s, followed by the presentation of a 2.5-s melody while a fixation cross remained on the screen. Each trial ended with a 5.5-s silent interval before the onset of the next trial. In total, 120 trials were presented in randomized order (60 listen, 60 memorize) and each melody was repeated 20 times. Participants were randomly assigned to one of two counterbalanced groups. Group 1 (n = 33) passively listened to melodies s1, m1, and c1 and actively memorized s2, m2, and c2, whereas Group 2 (n = 34) performed the opposite assignment.

After the MEG session, participants took a short break before proceeding with the behavioral part of the study. This included an old/new recognition task (Bonetti et al., 2021; Bonetti et al., 2023; Bonetti et al., 2024a; Bonetti et al., 2024b; Bonetti et al., 2024c; Bonetti et al., 2025; Fernández-Rubio et al., 2022a; Fernández-Rubio et al., 2022b; Fernández-Rubio et al., 2024; Nartallo-Kaluarachchi et al., 2025; Serra et al., 2023), complexity and pleasantness ratings of each melody, a background questionnaire, and the Goldsmiths Musical Sophistication Index (Müllensiefen et al., 2014).

During the old/new recognition task, participants were presented with the same melodies from the encoding task and with variations of these. For each melody, participants were requested to press one button if they had heard the melody (memorized) and a different button if they had not heard the melody before (novel) (**Figure 1b**). The task was completed on site at Aarhus University Hospital (AUH) on the same day as the MEG recording, one day later, and 10 days later using the Cognition platform (www.cognition.run).

Structural MRI data were acquired in a separate session following the MEG and behavioral testing.

### Data acquisition

MEG data were acquired in a magnetically shielded room at AUH using a 306-channel TRIUX MEG system (Elekta Neuromag, Helsinki, Finland). Data were acquired at a sampling rate of 1000 Hz with an analogue filtering of 0.1–330 Hz. Participants’ head shapes and the positions of four Head Position Indicator (HPI) coils were digitized with a 3D tracking system (Polhemus Fastrak, Colchester, VT, USA). HPI coils monitored head position throughout the recording and were later used for movement correction. Bipolar electrodes were placed to record electrooculography (EOG) and electrocardiography (ECG) signals for subsequent artifact rejection.

MRI data were acquired in a separate session on a CE-approved 3T Siemens scanner at AUH. Structural T1-weighted images (MPRAGE with fat saturation) with a spatial resolution of 1.0 × 1.0 × 1.0 mm were obtained using the following parameters: echo time (TE) = 2.61 ms, repetition time (TR) = 2300 ms, reconstructed matrix = 256 × 256, echo spacing = 7.6 ms, and bandwidth = 290 Hz/px.

### MEG data preprocessing

Raw MEG sensor data from 204 planar gradiometers and 102 magnetometers were first preprocessed using MaxFilter (v2.2.15) (Taulu & Simola, 2006) to suppress external interference. The following parameters were applied: spatiotemporal signal space separation (SSS), downsampling from 1000 Hz to 250 Hz, and movement compensation based on HPI coil signals (default step size = 10 ms). The correlation limit between the inner and outer subspaces used to reject overlapping signals during spatiotemporal SSS was set to 0.98.

The data were then converted to Statistical Parametric Mapping (SPM) format and further preprocessed and analyzed in MATLAB (MathWorks, Natick, MA, USA) using a combination of in-house scripts (https://github.com/leonardob92/LBPD-1.0.git) and the Oxford Centre for Human Brain Activity (OHBA) Software Library (OSL) (Woolrich et al., 2011), which builds on FieldTrip (Oostenveld et al., 2011), FSL (Woolrich et al., 2009), and SPM12 (Penny et al., 2011). Continuous data were visually inspected with OSLview to identify and remove artifacts. Independent component analysis (ICA) was used to discard components related to eye movements and cardiac activity (OSL implementation).

Finally, the continuous data were epoched into 120 trials (60 listen, 60 memorize) and baseline-corrected by subtracting the mean signal in the pre-stimulus baseline from each post-stimulus epoch. Each epoch lasted 3.1 s (0.1-s baseline, 2.45-s stimulus, 0.55 s silence). To focus on differences in encoding, and in line with our previous study that compared tonal and atonal melodies during recognition (Fernández-Rubio et al., 2022b), only “memorize” trials for tonal and atonal melodies were included in further analyses.

### Source reconstruction and regions of interest

Using the information collected with the 3D digitizer, the MEG sensor data and individual MRI T1-weighted images were co-registered. A single-shell forward model using an 8-mm grid and beamforming algorithm as inverse solution were employed (see Bonetti et al. (2024b) for details). This procedure returned a time series for each of the 3559 brain sources (**Figure 1c**).

Building on our previous studies of musical memory recognition (Bonetti et al., 2024b; Bonetti et al., 2024c), we analyzed a set of anatomically defined regions of interest (ROIs): the left and right Heschl’s gyrus, left and right hippocampus, medial cingulate gyrus, and anterior cingulate gyrus. Using the automated anatomical labelling (AAL) parcellation method (Tzourio-Mazoyer et al., 2002), we extracted a time series for each of these six ROIs (**Figure 1d**). For each ROI, we identified the corresponding brain voxels and averaged their time series. To resolve the sign ambiguity of the evoked response time series, we aligned each ROI’s sign with the N100 response to the first tone of the melodies (Bonetti et al., 2024b; Bonetti et al., 2024c; Fernández-Rubio et al., 2022b). This procedure yielded a final time series for each ROI and experimental condition (**Figure 1e**).

### Statistical analysis

Statistical analyses were conducted to compare the time series of tonal versus atonal melodies at each ROI. For each timepoint and ROI, two-sided t-tests were computed, and corrections for multiple comparisons were applied using a cluster-based permutation test (Maris & Oostenveld, 2007). This analysis consisted of computing two-sided t-tests independently for each timepoint and identifying clusters of neighboring significant timepoints by applying a threshold to the statistical test results (α = 0.05). Next, 1000 permutations were performed, shuffling the labels of the two experimental conditions for each participant and recalculating the statistics. For each permutation, the maximum cluster size of significant timepoints was computed, yielding a reference distribution of significant clusters. Clusters in the original data were deemed significant only if their sizes exceeded the maximum cluster size observed in 99.9% of the permuted data.

To compare participants’ perceived complexity of the two melody types, we used a Wilcoxon rank-sum test on the complexity ratings. This non-parametric test was selected due to the ordinal nature of the data (scores on a 7-point Likert scale). The analysis was conducted using the *wilcox.test* function from the *stats* package in R (version 2024.12.1+563), with statistical significance set at p < 0.05.

To model the effect of melody complexity on response accuracy over time in the old/new recognition task, we fitted a generalized linear mixed-effects model (GLMM) with a binomial error distribution and logit link function using the *glmer* function from the *lme4* package in R. This approach was chosen because the response data were discrete distributions indicating the probability of giving a correct response (*n*) out of the total number of responses (*k*). Fixed effects included melody complexity (simple, complex), time (session 1, session 2, session 3), and their interaction, while participant was included as a random effect. Post hoc pairwise comparisons were conducted using estimated marginal means (function *emmeans* from the *emmeans* package), with p-values adjusted using the Tukey method for multiple comparisons.

## ACKNOWLEDGEMENTS

The Center for Music in the Brain (MIB) is funded by the Danish National Research Foundation (project number DNRF117), The Lundbeck Foundation (R469-2024-1573), and Købmand Herman Sallings Fond. L.B. is supported by Sapere Aude: Independent Research Fund Denmark (DFF) Research Leader (grant ID: 10.46540/5253-00003B), Lundbeck Foundation (Talent Prize 2022), Carlsberg Foundation (CF20-0239), Center for Music in the Brain, Linacre College of the University of Oxford, and Nordic Mensa Fund. M.L.K is supported by Center for Music in the Brain and Centre for Eudaimonia and Human Flourishing, which is funded by the Pettit and Carlsberg Foundations. We thank Nikita Joe, Shiraz Ben-Shoshan, and Samuel William Nehrer for their assistance with data collection.

## AUTHOR CONTRIBUTIONS

G.F.-R., L.B., and E.B. designed the study. L.B., M.L.K., and P.V. recruited the resources for the experiment. G.F.-R. collected the data. G.F.-R. and L.B. performed pre-processing, source reconstruction, and statistical analyses. G.F.-R. and L.B. prepared the figures and wrote the first draft of the manuscript. All the authors contributed to and approved the final version of the manuscript.

## CODE AVAILABILITY

The main data analysis pipeline used in this study is available at the following link: https://github.com/gemmaferu/encoding-complexity

## SUPPLEMENTARY MATERIAL

### Supplementary figures

**Figure S1.**
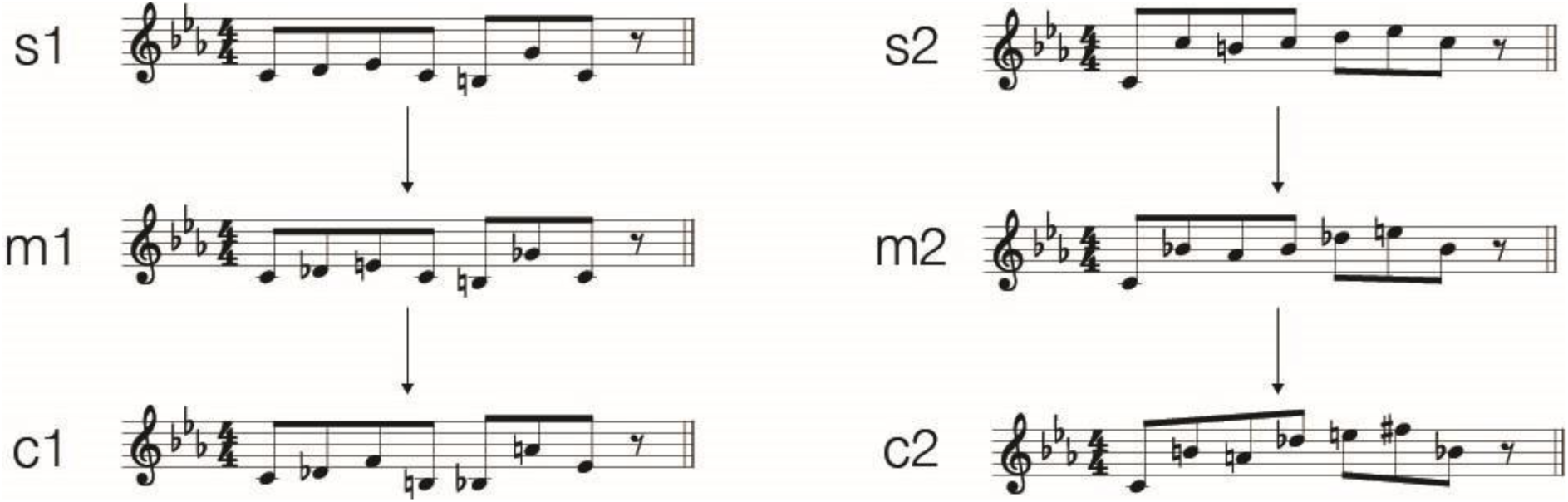
Musical stimuli. The stimulus set comprised two tonal (s1, s2), two semi-tonal (m1, m2), and two atonal melodies (c1, c2), matched for rhythm, timbre, tempo, and meter.

### Supplementary tables

**Table S1.** Statistical analysis of source-localized brain activity. Results of statistical analyses conducted independently for six regions of interest (ROIs). At each timepoint, t-tests compared simple versus complex conditions. Multiple comparisons were corrected using cluster-based permutations. Significant clusters are reported for each ROI, including cluster size, temporal extent of the cluster, p-value, and peak t-value.

| Left Heschl's gyrus | Cluster size | Start (s) | End (s) | p-value | Max t-value |
| --- | --- | --- | --- | --- | --- |
|  | 11 | 0,984 | 1,024 | < 0,001 | 3,527 |
|  | 8 | 2,352 | 2,380 | < 0,001 | 3,139 |
| Right Heschl's gyrus | 7 | 0,864 | 0,888 | < 0,001 | 3,421 |
|  | 5 | 1,536 | 1,552 | < 0,001 | -3,806 |
| Left hippocampus | 6 | 0,868 | 0,888 | < 0,001 | -2,359 |
|  | 11 | 1,244 | 1,284 | < 0,001 | -3,069 |
|  | 9 | 1,324 | 1,356 | < 0,001 | -3,404 |
|  | 9 | 1,400 | 1,432 | < 0,001 | -3,269 |
| Right hippocampus | - | - | - | - | - |
| Medial cingulate gyrus | 5 | 0,504 | 0,520 | < 0,001 | 2,609 |
|  | 7 | 1,212 | 1,236 | < 0,001 | 3,218 |
| Anterior cingulate gyrus | 8 | 1,320 | 1,348 | < 0,001 | -2,715 |
|  | 6 | 1,416 | 1,436 | < 0,001 | -2,895 |
|  | 10 | 2,396 | 2,432 | < 0,001 | -3,755 |

